# 3D bioprinting of engineered living materials in support slurries for complex free-standing structures

**DOI:** 10.64898/2026.05.20.726626

**Authors:** Ram S. Gona, Hongyi Cai, Marie Olland, Manasi Subhash Gangan, Danielle T. Bennett, Urvi O. Mehta, Meredith N. Silberstein, Anne S. Meyer

## Abstract

The combination of synthetic biology and additive manufacturing has driven major changes in production of biomaterials, especially through the use of three-dimensional (3D) bioprinting to create engineered living materials. However, current fabrication methods can be limited by prohibitive hardware costs and the inability to maintain structural fidelity in complex, free-form living architectures. This work demonstrates how to build a low-cost, open-source 3D bioprinting platform that can make complicated bacterial structures with complex geometry and high dimensional accuracy. A commercially available, conventional fused deposition modeling 3D printer was modified to create a bioprinting system that is simple to build. The modified bioprinter, which costs around $450, is less expensive than many commercial bioprinters. This 3D-printing technology uses slurry-based support bath methods featuring low-cost gelatin and agarose microparticles, resulting in structures with a high aspect ratio (>8:1) and feature sizes as small as 260 μm. The optimization of critical printing settings, including the ability of the bioink to retract during non-print movements, resulted in a reduction of unwanted bacterial deposition by nearly two orders of magnitude. Long-term viability experiments showed that bacteria in the bioprints could survive for at least 28 days with nutrient supplementation. Additionally, 3D-printed engineered biofilms revealed that incubation conditions and extracellular matrix composition significantly impacted the mechanical properties of printed constructs, with tradeoffs between matrix production and mechanical integrity. This study showcases an accessible 3D bioprinting platform for advanced bioprinting technologies, enabling development of engineered living materials with potential applications in synthetic biology, biotechnology, and tissue engineering.

## Introduction

The intersection of synthetic biology and advanced manufacturing has resulted in significant transformations in the production of biomaterials, enabling the development of functional constructs with exceptional complexity, precise spatial organization, and inherent biological activity^1, 2^. This interdisciplinary field combining synthetic biology, biomaterials science, and 3D bioprinting, has undergone significant advancements via the design and fabrication of engineered living materials (ELMs) and complex tissue-like constructs^3, 4^. Engineered living materials (ELMs) are characterized by their use of living cells or biologically synthesized biomaterials as essential elements. These materials can demonstrate unique abilities to sense, respond to, adapt to, and potentially modify their surroundings. These characteristics allow ELMs to be applied as adaptive or self-healing materials, environmental biosensors, advanced therapeutic delivery systems, or essential tools for investigating complex biological processes, including biofilm formation^5, 6^. In addition, the ongoing advancement of 3D bioprinting technologies offers the ability to control the spatial accuracy of the deposition of biomaterials, cell-loaded bio-inks, and living biological agents^7^.

The fundamental goal of ELM research lies in harnessing the intrinsic capabilities of organisms, especially microbes, to generate biomaterials with desired characteristics. Bacteria can produce an extensive variety of functional biomolecules and exhibit complex self-assembly behaviors, leading to the establishment of multicellular communities known as biofilms^8, 9^. Biofilms are defined by the production of a self-generated extracellular matrix (ECM), which consists of polysaccharides, proteins (including functional amyloids such as curli), lipids, and extracellular DNA. These matrix components provide structural integrity and protect the biofilm community from external stresses^10, 11^. Bacterial cellulose is a biopolymer synthesized by bacteria that serves as a primary structural component of the ECM for many types of biofilms. High-purity bacterial synthesis of bacterial cellulose occurs in specific strains, such as *Acetobacter xylinum* and naturally biofilm-forming or genetically modified *Escherichia coli* strains. Bacterial cellulose exhibits numerous advantageous materials characteristics, including a high mechanical strength, significant water retention, natural biocompatibility, and biodegradability^12, 13^, which can improve the materials properties of ELMs^14, 15^.

Additive manufacturing, particularly three-dimensional (3D) bioprinting, is an essential tool for the production of ELMs. This approach enables the accurate placement of cell-laden bio-inks and functional biological agents into spatially ordered structures^7, 16, 17^. Extrusion-based bioprinting is a technique frequently utilized to accurately deposit continuous bio-ink through a micro-nozzle, allowing the layer-by-layer construction of complex 3D structures^18^. Hydrogels serve as the principal nutritional medium and scaffold material in the 3D bioprinting of ELMs^19, 20^. They consist of three-dimensional networks of hydrophilic polymers that can absorb and hold significant amounts of water, thus creating a biocompatible and cell-friendly environment^21^. Alginate, a polysaccharide derived from brown seaweed, and gelatin, a denatured collagen, are naturally-derived hydrogels that are extensively utilized in bioprinting applications due to their biocompatibility and favorable rheological properties^22, 23^. Despite many advancements, the printing of intricate or delicate structures using hydrogel-based bio-inks continues to be a significant challenge. The inherent low viscosity and poor mechanical strength of most bio-inks can result in printed structures collapsing under their own weight, thereby preventing the fabrication of complex geometries such as overhangs, internal voids, or high-aspect-ratio objects. Consequently, direct bioprinting methods are often limited to the creation of simple, stacked geometries or structures of limited height^24-28^. While recent advancements in light-based stereolithography have enabled the fabrication of intricate structures, these approaches often require specialized resin chemistries and can be prohibitively expensive for bacterial research^29,30^.

To mitigate the limitations of hydrogel-based bio-inks, researchers have developed innovative strategies, such as using temporary sacrificial support materials and utilizing embedded 3D bioprinting processes^31, 32^. One example is the Freeform Reversible Embedding of Suspended Hydrogels (FRESH) technology, initially used for the printing of soft materials such as collagen and alginate to fabricate mammalian tissue constructs^33^. This technique enables the creation of intricate biological structures by extruding bio-ink into a support medium that functions as a yield-stress fluid. The use of support baths can prevent the collapsing of printed structures due to gravity, allowing the application of lower-viscosity bio-inks in the fabrication of large three-dimensional structures^34, 35^. Recently, this approach has been expanded to microbial systems by using silicone-based granular gels as a support bath^36^. However, non-reversible supports can complicate the extraction of delicate hydrogel constructs.

Although significant advancements have been made in 3D bioprinting, the high costs of commercial bioprinters can limit the accessibility of this technology for many research groups. To address this, several groups have demonstrated that low-cost, commercially available fused deposition modeling (FDM) printers can be modified into functional bioprinters for patterning bacteria^16, 26, 28^. However, these accessible do-it-yourself (DIY) systems have predominantly been limited to direct printing onto agar or other flat surfaces, with the resultant geometric constraints of that method. A critical gap has remained: the adaptation of these affordable, open-source platforms for use with advanced support baths to enable the fabrication of complex, free-standing ELMs. This work addresses the gap by introducing a low-cost, open-source 3D bioprinting platform optimized for printing in slurry-based support baths. We demonstrate the modification of a commercial FDM printer to create an accessible and high-performance platform for bacterial bioprinting. Customized agarose and gelatin support slurries were developed that enable the printing of engineered *Escherichia coli* bacteria integrated within an alginate-based bio-ink into intricate, self-supporting structures (Figure 1). Optimization of material retraction during non-printing motion of the printing nozzle resulted in improved printing resolution. This method can generate high-aspect-ratio structures with high fidelity and can sustain prolonged survival of embedded bacteria. Finally, we demonstrate that incubation conditions can regulate the accumulation of extracellular matrix (ECM) components and concomitantly tune the elastic modulus and ultimate tensile strength of the bioprints. Ultimately, this low-cost, open-source platform can democratize the fabrication of complex living architectures, with the potential to accelerate innovation in the field of engineered living materials.

**Figure 1.**
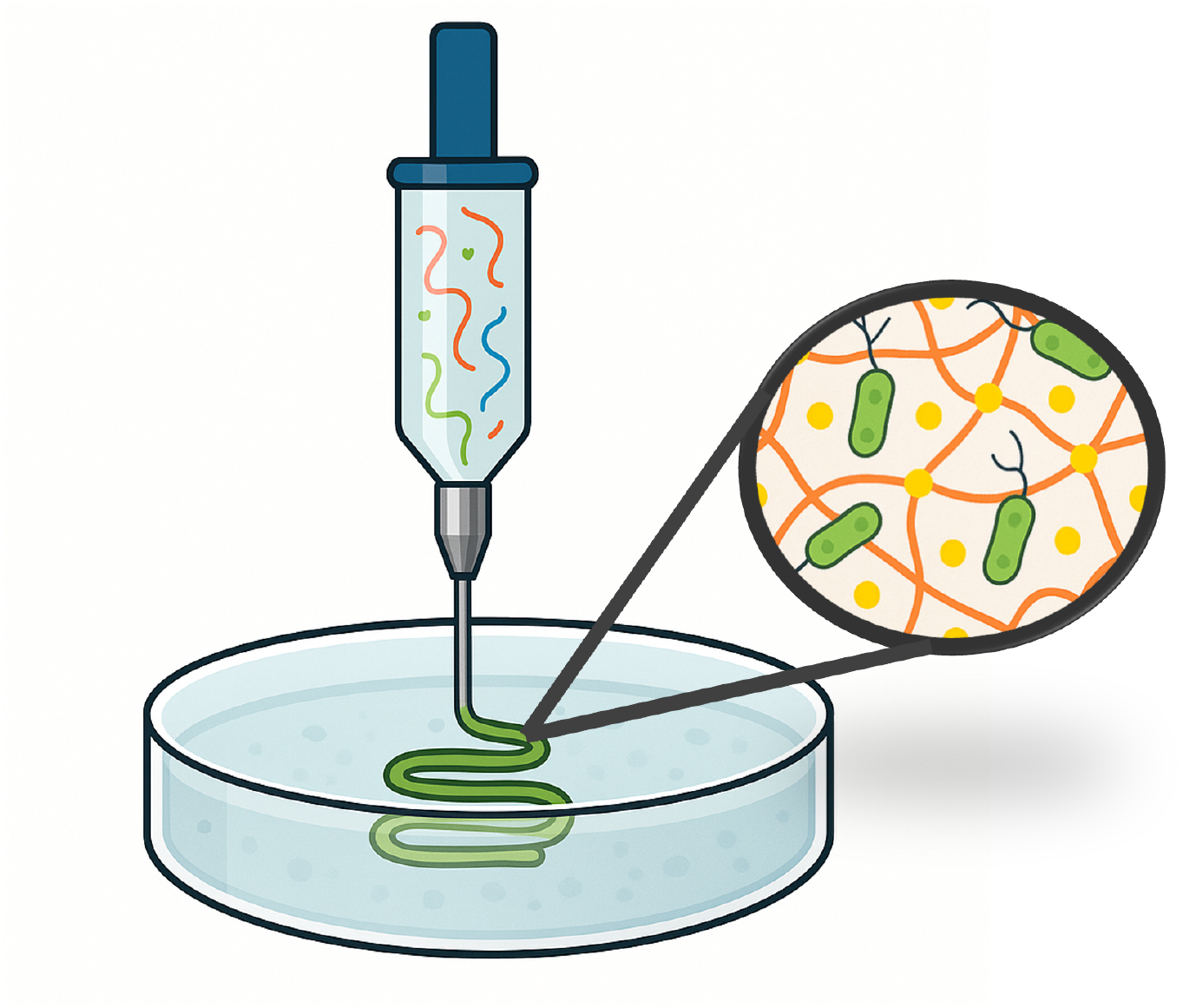
Graphical representation of the slurry-based 3D-bioprinting process. The syringe nozzle extrudes a bacterial bio-ink directly into a support slurry bath. The slurry is enriched with calcium ions (Ca^2+^). When the sodium alginate bio-ink is deposited, the divalent Ca^2+^ ions crosslink the alginate strands, forming a stable hydrogel matrix that solidifies the printed structure and encapsulates the living bacteria.

## Results

### Development of an inexpensive, open-source 3D bioprinter

To help advance the widespread adoption of 3D bioprinting technology in research settings, we developed a cost-effective, open-source 3D bioprinter by modifying a commercial fused deposition modeling printer (Creality Ender 3, ∼$450) (Figure 2A). This platform lowers the financial barrier to entry relative to commercial bioprinters, which generally have costs of thousands of dollars, without compromising the ability to generate high-resolution prints. To adapt the device for bioprinting, the conventional FDM print head and heating element were removed and substituted with a custom-engineered syringe pump assembly for extrusion of bioink (Figure 2B). The syringe pump design drew inspiration from the open-source Poseidon syringe pump system^37^, which required significant modifications to enable compatibility with the Creality printer platform while preserving its structural integrity and precision. Since the pump would be mounted onto the printer’s moving carriage, it needed to be lightweight to prevent excessive stress on the printer motors that drive it. The structural components of the pump were optimized by eliminating material from non-critical regions. The use of 90% infill when 3D-printing pump components allowed for mechanical stability to be retained. The optimization on the syringe body achieved 29% reduction in overall weight compared to the original open-source design. Custom mounting brackets were developed that are compatible with the Ender 3 extruder carriage, allowing for direct attachment of the syringe pump to the printer’s X-axis. The pump’s plunger mechanism was modified to wrap around both the top and bottom edges of the syringe’s plunger. This design enabled both extrusion and retraction of bio-ink, to allow for more precise control of bio-ink deposition and for a reduction in unintended material flow during printhead movements.

**Figure 2.**
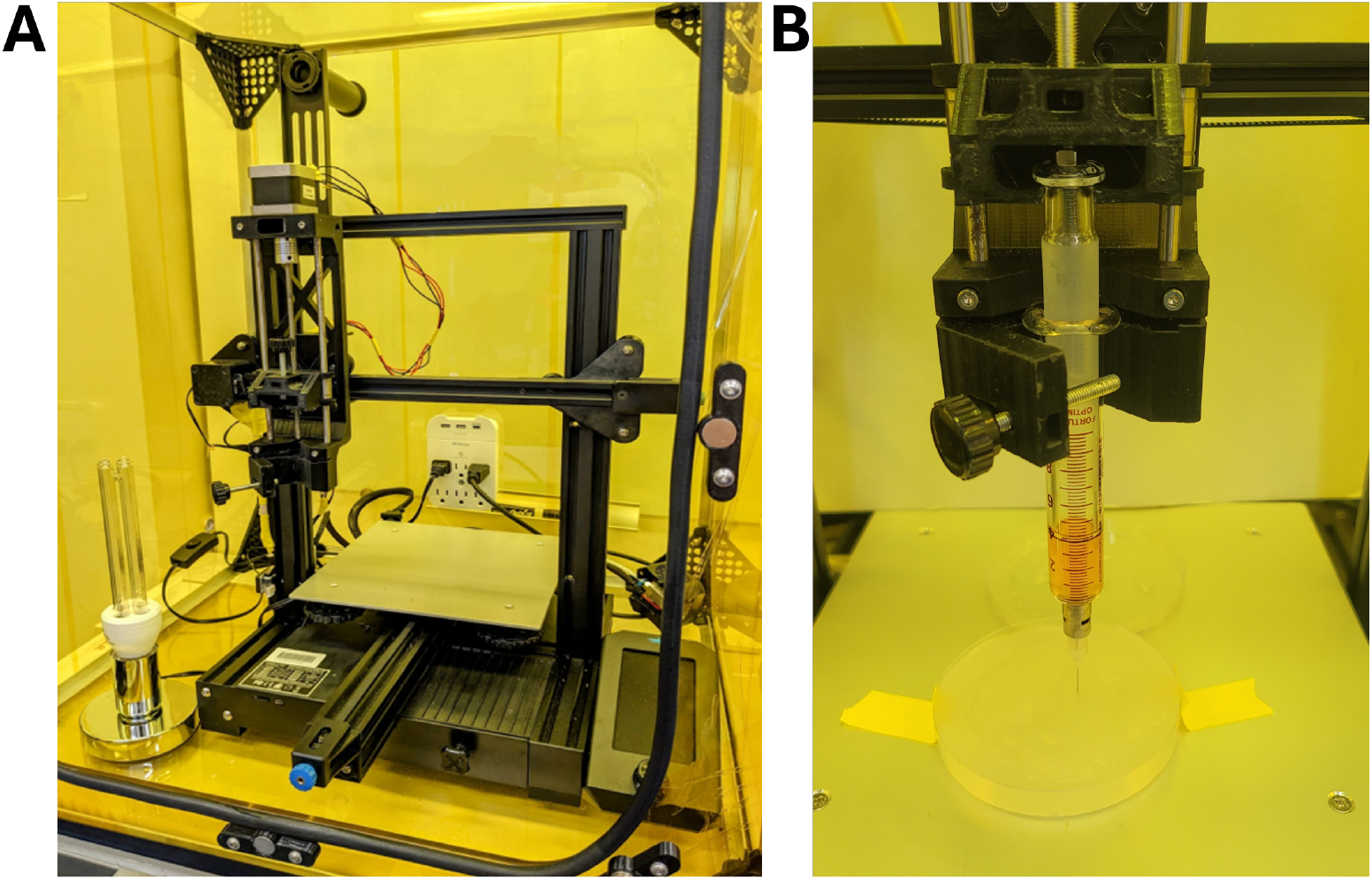
3D bioprinter setup for bacterial printing. (A) DIY 3D bioprinter inside a UV-blocking, polyimide-coated polycarbonate enclosure that confines BSL-2 organisms and prevents escape of 254 nm UVC sterilization light. (B) The custom syringe extruder: a 5 mL Luer-lock syringe driven by a DIY syringe pump replaced the original filament head, allowing precise extrusion (↓) and fast retraction (↑) of the bio-ink to prevent stringing. Printing occurs in a 90 mm Petri dish containing 1% (w/v) agarose slurry.

A major improvement in our design was the implementation of glass syringes instead of traditional plastic syringes^16, 38^. This modification enhanced printing resolution by eliminating the mechanical compliance associated with plastic syringes, which can result in variable material flow during printing processes. The glass syringe assembly was secured into the syringe pump with custom-designed clamps that ensured syringe alignment and prevented movement during printing processes. The modified printer was enclosed within a customized polycarbonate enclosure to enable work with Biosafety Level 2 (BSL-2) organisms if desired and to maintain sterile printing conditions (Figure 2A). A UVC lamp (254 nm) was integrated for surface sterilization between printing sessions. Additionally, temperature and humidity sensors were included to monitor the local environmental conditions during the printing process. The alterations enabled the printer to operate in conditions that promoted both sterility and the viability of the printed organisms.

### Optimization of G-code generation and bioprinting parameters

To adapt the printer for hydrogel extrusion and generate G-code instructions, several key parameters were modified in the open-source slicing software (Slic3r). The nozzle diameter was set at 0.254 mm to correspond with the inner diameter of the 26-gauge dispensing needle utilized in the bioprinter. The extrusion multiplier, which calibrates the software-entered flow rate value to the actual flow rate, required extensive modification as the bioprinter extrudes bio-ink instead of flowing plastic filament. Experimental testing revealed that an extrusion multiplier of 0.004 resulted in a flow rate of approximately 2 mL/hour at a print speed of 23 mm/s. These parameters were applied for printing the alginate-based bio-inks utilized in our experiments, since they allowed for optimal shape fidelity, minimizing defects such as bio-ink bulging via over-extrusion, or tearing via under-extrusion. During printing, bio-ink was observed to seep out of the dispensing needle when the extrusion was halted to reposition the printhead, due to the high viscosity of the bio-ink. In order to mitigate undesired material deposition during these non-printing movements, the syringe pump retraction parameter was chosen. Experiments indicated that a retraction distance of 0.17 mm yielded minimal off-target placement of alginate-based bioinks during bioprinting, which was optimized by evaluating the amount of bio-ink that oozed out at different retraction values. The shorter retraction distance, in contrast to the standard 3-5 mm used in FDM printing, indicates the distinct rheological properties of hydrogels versus thermoplastics^39^.

### Slurry-based support baths enable complex free-standing bacterial structures

Direct printing onto agar or other solid substrates has been shown in prior bacterial bioprinting methods^16, 17, 26, 28^; however, this approach presents limitations regarding the structural complexity and vertical dimensions of the printed constructs. To address these limitations and enable the fabrication of intricate, high-aspect-ratio bacterial structures, we modified and developed two types of slurry-based support bath techniques: gelatin slurry and agarose slurry. The slurries were created by casting agarose or gelatin gels and then blending them with a low-cost kitchen blender to create microparticles roughly 25-35 μm in diameter (Supplementary Figure 1). The slurries demonstrated yield-stress fluid properties, flowing as a liquid when a specific minimum shear stress is exerted by the moving needle during the printing process but acting as a solid in the absence of external stress to provide support to the printed structure post-extrusion^30,37^. Preparation of the gels by dissolving the initial agarose or gelatin powder in a solution of CaCl_2_ allowed for the resultant slurry microparticles to deliver Ca^2+^ ions to the bioink upon extrusion at any location within the slurry, cross-linking the alginate polymers of the bio-ink to form a polymerized hydrogel.

The gelatin slurry method was inspired by the Freeform Reversible Embedding of Suspended Hydrogels (FRESH) technique^33^, originally developed for bioprinting mammalian tissue. The gelatin slurry composition was optimized for bacterial bioprinting by adjusting the gelatin concentration to 4.5% (w/v) and the CaCl_2_ concentration to 11 mM. This composition provided sufficient structural support for extruded bio-ink and allowed efficient crosslinking of the alginate bio-ink upon deposition, resulting in a stable hydrogel structure. The low melting point of the gelatin slurry offered the benefit of enabling the extraction of the printed construct after 3D bioprinting. Heating the entire bath to 40°C caused the thermoreversible gelatin to melt, allowing for the removal of the crosslinked alginate structure containing the embedded bacteria (Figure 3). The relatively low temperature and short duration of heating required to melt the gelatin slurry, compared to other hydrogels typically used for support slurries^36, 40^ that have higher melting points, reduces adverse effects on the embedded bacteria

**Figure 3.**
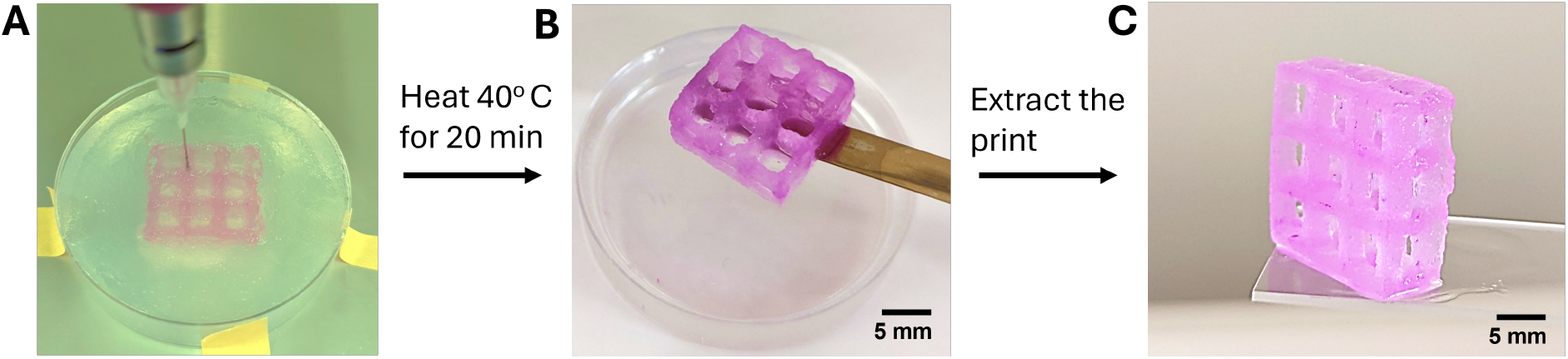
Gelatin slurry-based support baths enable the printing and extraction of complex free-standing structures. (A) A complex lattice structure was 3D printed into a gelatin slurry support bath. (B) The gelatin support was melted by heating the bath to 40°C for 20 minutes, allowing for the extraction of the printed construct. (C) The final extracted bioprint was free-standing, demonstrating its complex geometry and structural integrity even after removal from the support bath.

An alternative approach, inspired by the work of Mirdamadi et al.^40^, utilizes agarose microparticles for applications requiring extended incubation within the support medium. Agarose slurry was prepared using 1% (w/v) agarose dissolved in a solution of 11 mM CaCl_2_. This process offered mechanical support and rapid bio-ink gelification similar to that of the gelatin slurry. Since agarose can maintain stable polymerization at temperatures up to 95°C, this slurry could be used as an incubation medium providing support for the bacteria-embedded structures after the printing process.

Both slurry baths allowed the formation of complex bacterial structures. To demonstrate the capabilities of our modified open-source bioprinter and support slurry printing technique, a three-dimensional lattice structure was designed using computer-assisted design software (Figure 4A). This design was printed within the support slurries to create structures with heights surpassing 20 mm (Figure 4B). Once removed from the slurry, the lattice was able to be free-standing while retaining its geometrical features, including overhangs, internal channels, and hollow structures (Figure 4C-D), which are difficult to achieve with traditional bioprinting methods that entail direct printing onto solid surfaces. The complex three-dimensional structures that could be fabricated using this slurry-based approach represent an improvement over earlier bacterial bioprinting techniques, which primarily focused on two-dimensional or limited three-dimensional configurations^16, 24, 31^.

**Figure 4.**
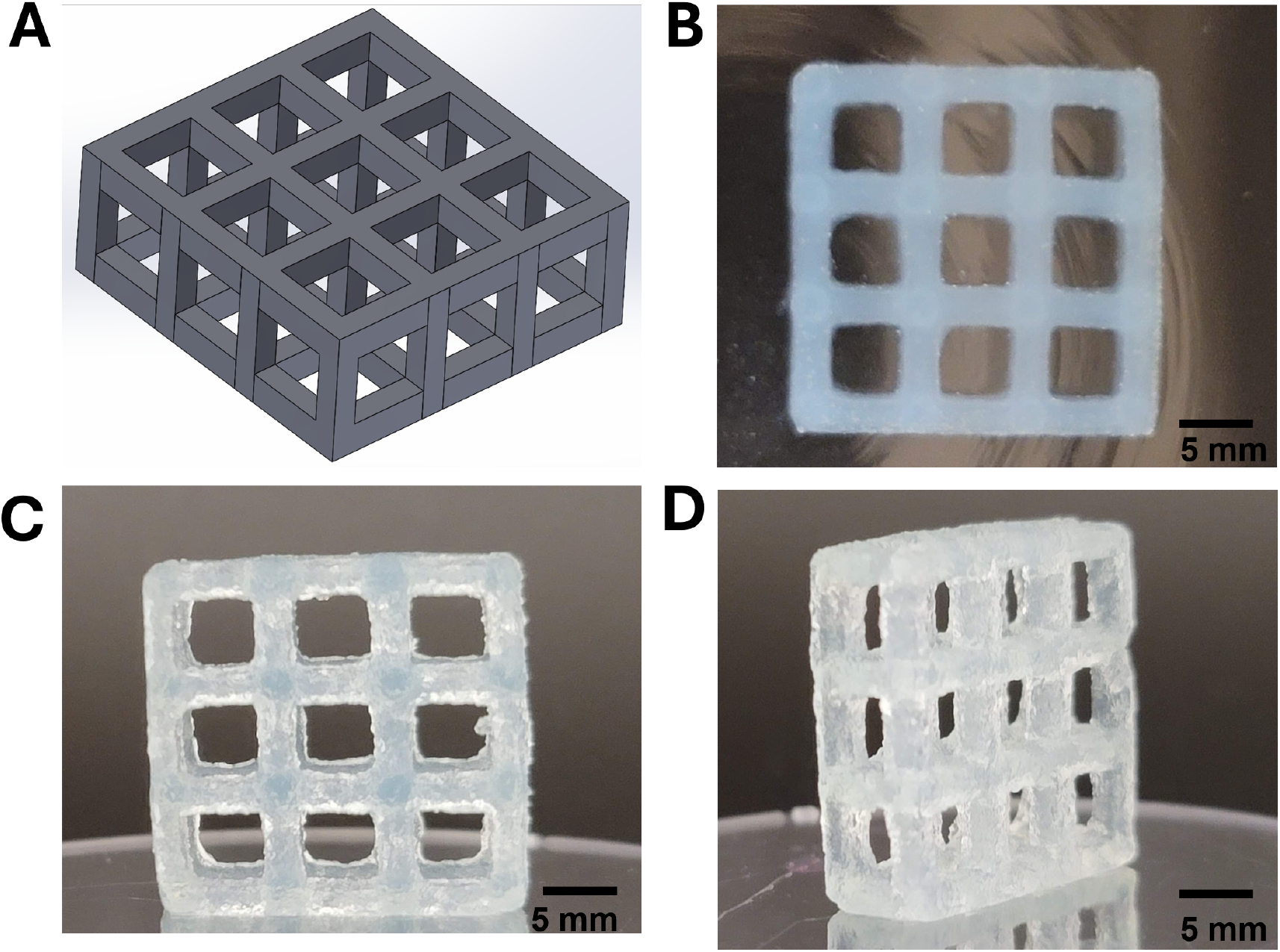
Free-standing 3D-printed square lattice structure composed of alginate hydrogel laden with meffBlue-expressing *E. coli*^41^. (A) Computer-aided design (CAD) model of a lattice structure illustrating the presence of complex hollow features. (B) Top view of the square lattice structure 3D printed with optimized parameters in the melted gelatin slurry before its extraction. (C) and (D) Top and isometric view of the 3D-printed structure removed from the slurry bath. The structure measured 21 mm in height and length and 6.5 mm in width, with the center hollow squares measuring 5 mm in length.

### Retraction capability reduces off-target bacterial placement

A major challenge in bioprinting living materials is attaining precise spatial control over the deposition of the bio-ink. During the printing of complex structures that have hollow void or disconnected features, the print nozzle needs to move throughout the slurry media, both when it is and when it is not actively printing. Leakage of bio-ink containing bacteria during these frequent movements will result in prints with blurred edges, misshapen features, or deposition of bio-ink in undesired locations. Experiments were performed to examine the influence of retraction capability, which is the ability to reverse the extruder motor direction to minimize bio-ink oozing during non-printing movements, on print quality and bacterial distribution.

In these experiments, a chromogenic *Escherichia coli* strain (Top10 pSB1C3-BBa_K1033923) was utilized, which expresses the spisPink chromoprotein extrachromosomally under the constitutive J23110 promoter^42^. The highly pigmented strain was suspended in 4% (w/v) alginate bio-ink and printed into hollow cylindrical structures within an agarose slurry support bath using the modified bioprinter. Each hollow cylinder was 10 mm in height, with a 10-mm external diameter and a 5-mm internal diameter. The print consisted of four identical, adjacent cylinders printed in the same print run, such that the printhead moved extensively between the cylinders through the slurry during the print process. A comparative analysis was conducted to evaluate the effect of retraction on print quality by examining identical print patterns produced with and without the retraction feature enabled (Figure 5A-B). When retraction was disabled (Figure 5A), bacterial streaking was visible between the cylindrical structures, resulting in unwanted connections that prevented the separation of individual cylinders. The streaks were observed to result from passive material flow during non-printing movements, a common issue in extrusion-based bioprinting systems^7, 24^. With retraction enabled (Figure 5B), the printed structures displayed cleaner boundaries and minimal visible contamination between structures.

**Figure 5.**
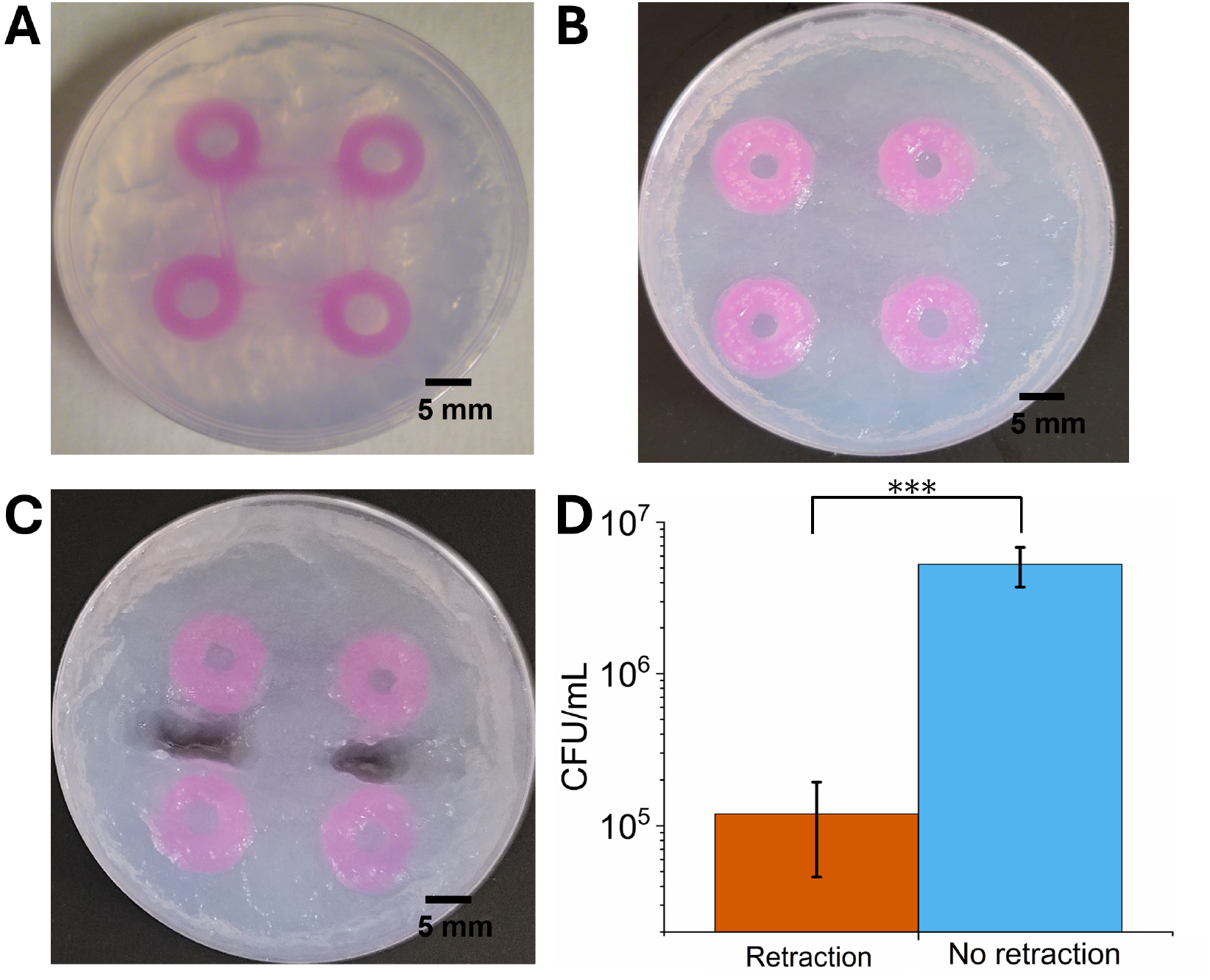
Retraction capability significantly reduces off-target bacterial placement between printed structures. (A) Representative image of cylindrical structures printed with retraction disabled, showing bio-ink streaks (pink) between intended features. (B) Identical print pattern produced with retraction enabled at 0.17 mm retraction distance, showing clean boundaries and minimal inter-structure contamination. (C) Illustration of the sampling method for quantitative analysis of bacterial contamination in the inter-structure regions. (D) CFU analysis of samples extracted from inter-structure regions, either with (orange) or without (blue) retraction capability enabled. Data represent mean ± SEM (n = 4); statistical significance determined by t-test (*** p < 0.001).

To quantitatively evaluate the effect of retraction on bacterial contamination between printed structures, samples of equal mass were taken from the inter-structure regions of the agarose slurry (Figure 5C). Colony-forming unit (CFU) analysis was conducted to determine the concentration of viable bacteria in the slurry samples. Prints generated without retraction exhibited 5.3 × 10^6^ CFU ml^−1^ in the inter-structure regions, while prints with retraction demonstrated approximately 10^5^ CFU ml^−1^ (Figure 5D), reflecting a statistically significant reduction of nearly two orders of magnitude. This optimized retraction feature mitigates a limitation of numerous DIY bioprinting systems, which frequently exhibit material oozing during non-printing movements. The decrease in undesired bacterial deposition indicates an improvement in the precise spatial control of bacterial populations, which is important for the development of functional biofilm-based materials with specific properties^20, 28^.

### Print fidelity in slurry-based bioprinting

Earlier versions of DIY bioprinting, in which bio-ink was extruded onto solid substrates were often restricted to simple, low-aspect ratio geometries^7^. For example, Lehner et al.^16^ quantified this limitation at approximately 14 layers for alginate-based bacterial bio-inks. To determine the dimensional accuracy of slurry-printed structures, a quantitative layer-by-layer analysis of print fidelity was performed on high-aspect-ratio constructs. To help assess the consistency of 3D printing, colored bio-ink containing spisPink-expressing *E. coli* was utilized for ease of visualization. The bio-ink was printed into single-walled hollow cylindrical shapes generated by a single continuous pass of the printhead, making a 1-perimeter loop. Separate samples were printed with total layer numbers of 1, 10, 20, 30, 40, and 100 within an agarose slurry, and measurements of the total vertical height and the width of the printed wall were taken (Figure 6A). The width of a 1-layer print was 0.260 ± 0.027 mm, which closely aligns with the internal diameter of the 26-gauge needle used in the printing. The width was observed to increase from 1-layer (0.260 ± 0.027 mm) to 100-layer prints (0.548 ± 0.150 mm), partially due to settling of the soft hydrogel structure over time. However, the print width remained stable as the structure grew at lower layer numbers, measuring 0.346 ± 0.020 mm at 10 layers and 0.338 ± 0.019 mm at 20 layers. In comparison, previous open-source bioprinter systems produced structures with line widths that were approximately 5 times wider for a 1-layer print and that more than doubled in width between 1- and 14-layer prints^16, 38^, creating markedly lower-resolution structures that exhibited substantially more width expansion with increased layer numbers. The reduced width expansion demonstrated here in slurry-based printing is likely due to reduced gelation time for stacked layers, relative to printing onto solid substrates where cross-linking agents diffuse up through previously printed layers to solidify later layers.

**Figure 6.**
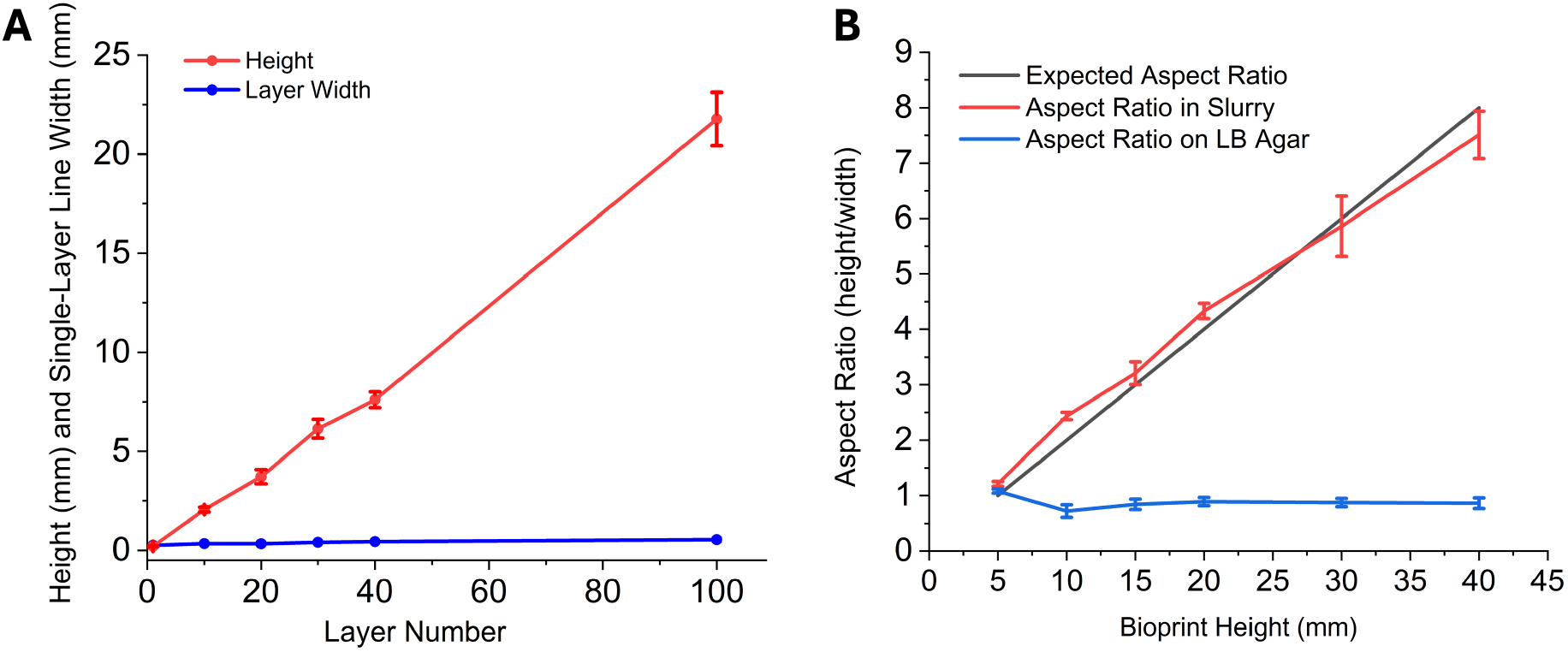
Slurry-based bioprinting enables high-aspect-ratio structures with precise dimensional control. (A) Analysis of the final measured height (red line) and line width (blue line) of bioprints containing spisPink-expressing *E. coli*, created with total layer numbers varying from 1-100. (B) The final achieved aspect ratios for cylindrical structures printed in an agarose slurry (red line) or directly onto an LB-agar surface (blue line) as well as the expected theoretical aspect ratios (black line), as a function of the expected bioprint heights. Error bars represent the standard deviation (n=3). The visibility of the printed structures was enhanced by a pink chromoprotein expressed by the bacteria.

The print height showed a linear increase corresponding to the number of layers printed, indicating deposition of consistent layers throughout the printing process. The height of a 1-layer print was measured at 0.260 ± 0.014 mm (Figure 6A), comparable to its width. For 100-layer prints, the total structure height was 21.8 ± 1.3 mm. The capability to print up to 100 layers while preserving reasonable dimensional accuracy marks a notable improvement over earlier DIY bioprinting methods^16, 38^. Slurry-printed structures were able to show more accurate material deposition due to the optimized extrusion mechanism and support by the slurry bath, preventing them from collapsing under their own weight after 15+ layers and resulting in more precise structures.

After establishing the maximum resolution of the print paths, the macroscopic structural fidelity of the modified bioprinter was evaluated by printing high-aspect-ratio structures. A comparative analysis was performed of prints generated in agarose slurry and those deposited directly onto agar substrates containing CaCl_2_ cross-linker. Colored bio-ink containing spisPink-expressing *E. coli* was printed into different cylindrical structures with a consistent expected outer diameter of 5 mm and expected heights ranging from 5-40 mm (approximately 25 to 200 layers). Structures printed directly onto Luria-Bertani (LB)-agar showed considerable structural collapse, showing little to no increase in aspect ratio with increasing heights (Figure 6B, Supplementary Figure 2). When printed in agarose slurry, the cylindrical structures preserved their intended shapes across all tested aspect ratios from 1:1 to 8:1 (Supplementary Figure 2). Quantification of the dimensions of the printed cylinders indicated that the shapes printed in the agarose slurry showed minimal deviation in aspect ratio from expected values, with the highest percent deviation of 6.3% measured for the 8:1 aspect-ratio print (Figure 6B), indicating that the slurry-based printing is able to maintain structural integrity in complex or challenging print forms.

### Long-term bacterial viability in slurry-printed constructs

The practical applications of bioprinted bacterial constructs depend on both their structural stability and the long-term viability of the embedded bacteria. To evaluate bacterial survival in agarose-slurry-printed constructs, a 28-day viability study was performed on spisPink-expressing *E. coli* 3D-printed into hollow cylindrical structures of 10-mm height, 10-mm outer diameter, and 5-mm inner diameter. Printed samples were incubated within the slurry at 37°C under two conditions: with LB nutrient supplementation and controls with added PBS buffer only. Prints were removed at regular intervals, and bacterial cell viability within the prints was assessed via colony-forming unit (CFU) assays. Cells in both experimental groups demonstrated an initial growth phase, with CFU per milliliter reaching a peak at 48-72 hours following printing (Figure 7). The LB-supplemented group attained a maximum concentration of 4.7 × 10^8^ CFU/mL at 72 hours, while the PBS control group reached its peak earlier at 48 hours with a concentration of 1.8 × 10^8^ CFU/mL. After the initial growth phase, both groups showed a decrease in viable cell counts, with a slower rate of decline for the LB-supplemented group. After 28 days of incubation, no viable colonies were observed in the PBS-incubated samples. In contrast, the LB-supplemented constructs exhibited a concentration of 6.25 × 10^5^ CFU/mL of viable bacteria at 28 days. The substantial increase in bacterial viability associated with nutrient supplementation can be attributed to the porous structure of the agarose microparticle network allowing the diffusion of nutrients to the embedded bacteria, thereby promoting their growth and metabolic activity of bacteria cells within the printed framework^43^. This result highlights the potential of slurry-based printing for creating durable engineered living materials.

**Figure 7.**
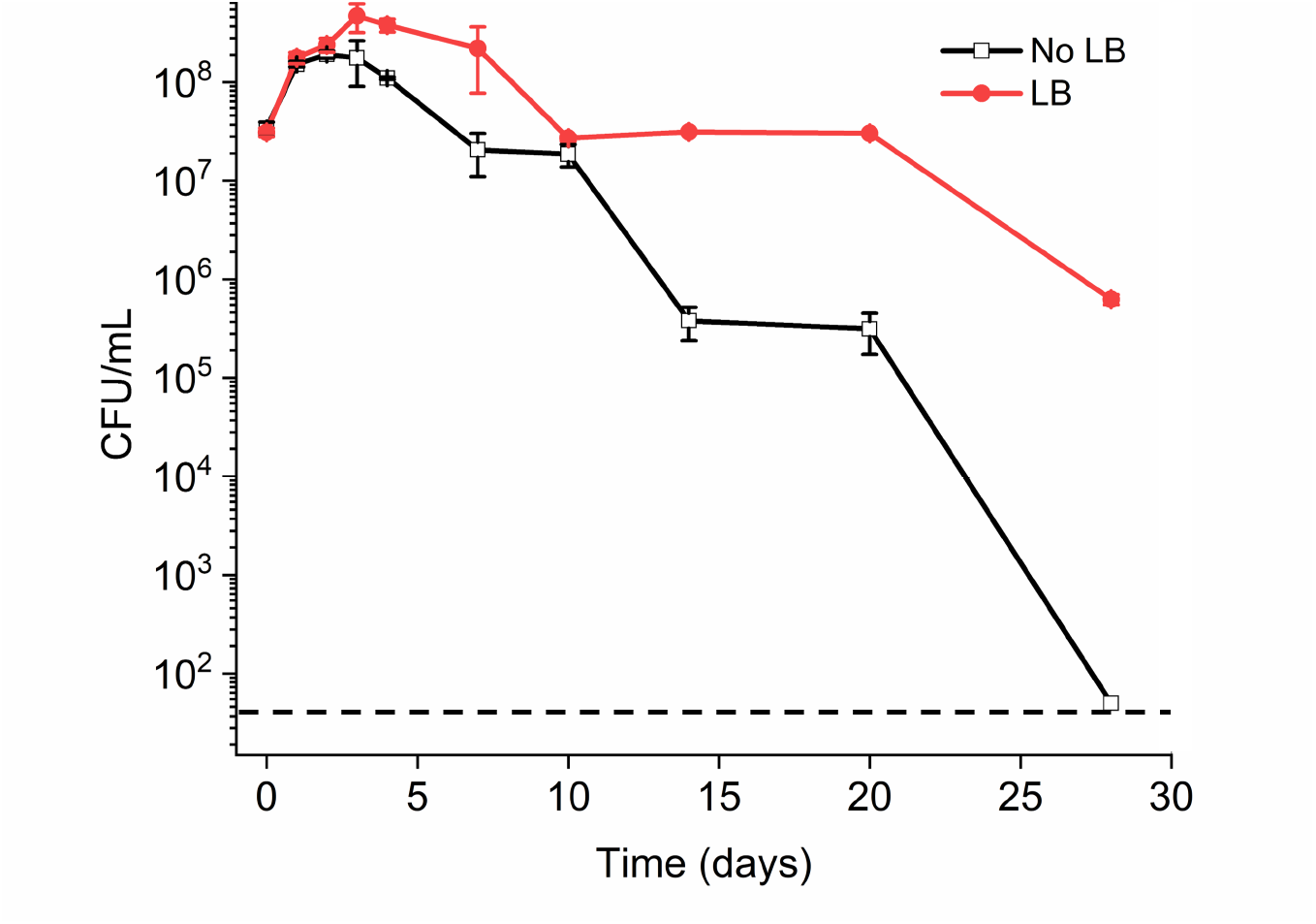
Nutrient supplementation enhances bacterial viability in printed constructs. The viability of *E. coli* in 3D-printed constructs was measured over a period of 28 days for constructs incubated with LB nutrients (red line, solid circles) and control constructs lacking LB supplementation (black line, open squares). The horizontal dashed line indicates the assay’s limit of detection. Data points are expressed as mean ± SEM (n=3).

### Incubation conditions of bioprints influence extracellular matrix production

The extracellular matrix (ECM) secreted by bacteria plays an important role in biofilm development. Additionally, it has been shown to impact mechanical stability and robustness of biofilms and alginate-based bioprints^20, 44^. To understand the impact of environmental variables on ECM formation in 3D-bioprinted bacterial constructs, the effect of incubation environment was examined. Three strains of *Escherichia coli* were tested: *E. coli* Nissle 1917 (EcN, curli^+^/cellulose^+^), a probiotic strain known for its production of both curli amyloid fibers and cellulose^45, 46^; *E. coli ΔcsgA* pCsgA-GFP (curli^+^/cellulose^−^), an engineered strain that expresses CsgA (curli) when induced but lacks cellulose^20^; and *E. coli ΔcsgA* pGFP (curli^−^/cellulose^−^), an engineered strain that produces neither curli nor cellulose^20^. Dog-bone-shaped structures were 3D printed within gelatin slurries using bio-inks containing these strains. The bioprints were extracted from the slurries and incubated for 7 days at 37°C under two different environmental conditions: one group was placed onto the surface of an LB-agar plate, providing exposure to air, while the other group was entirely submerged within liquid LB medium, resulting in a reduced-oxygen environment (Supplementary Figure 3).

After the incubation period, the ECM composition was evaluated within the 3D-printed structures. A Congo red assay was used to measure the total secreted amyloid polymers. Bioprints incorporating EcN (curli^+^/cellulose^+^) and *E. coli ΔcsgA* pCsgA-GFP (curli^+^/cellulose^−^) demonstrated significantly higher Congo red absorbance compared to control prints with *E. coli ΔcsgA* pGFP (curli^−^/cellulose^−^) for samples incubated on plates (Figure 8A). Incubation in liquid media significantly reduced the Congo red signal for both EcN and *E. coli ΔcsgA* pCsgA-GFP compared to their counterparts incubated on LB-agar (p < 0.001), and neither of these strains exhibited Congo Red signal that was significantly distinguishable from the non-curli-producing control (p > 0.05; Figure 8A). These data indicate that curli production by EcN and *E. coli ΔcsgA* pCsgA-GFP is more efficient in the oxygen-rich and dry environment of the agar plate than in the oxygen-limited and wet conditions of liquid media.

**Figure 8.**
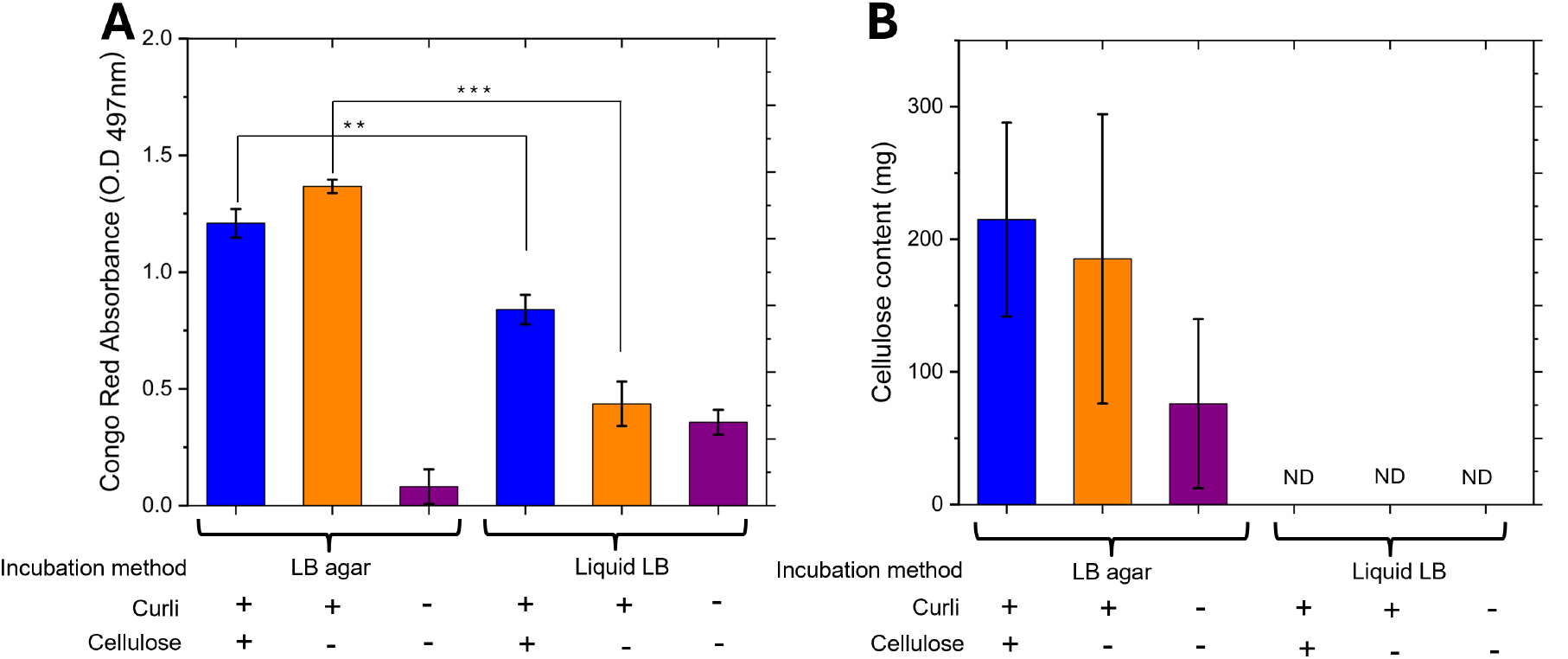
Extracellular matrix (ECM) production in 3D-printed *E. coli* constructs is influenced by incubation conditions. Dog-bone-shaped bioprints were incubated for 7 days at 37°C either on the surface of an LB-agar plate or fully submerged within liquid LB medium. (A) Congo Red adsorption and (B) bulk cellulose content measured by the anthrone assay for the bioprints. Bars represent mean ± SEM (n=3 or 4). Statistical significance was determined by a two-way ANOVA followed by Tukey’s post-hoc test. In Congo Red assays (A), asterisks indicate significant differences between marked groups (*p < 0.05; ** p < 0.01; *** p < 0.001); no significant differences were observed between strains in liquid media. In cellulose assays (B), no statistically significant differences were detected. ND indicates values below the detection limit of the assay.

An anthrone assay was conducted to quantify cellulose production in the incubated samples (Supplementary Figure 4). While bioprints with EcN incubated on an LB-agar surface exhibited the highest mean cellulose content among all groups, statistical analysis did not identify significant differences between strains or incubation conditions due to high variability across the replicates (Figure 8B). The cellulose values for bioprints submerged within liquid LB medium were reproducibly lower than those on plates and were at levels below the detection limit of the assay. This result is consistent with significantly enhanced rates of bacterial cellulose biosynthesis observed for enterobacteria under oxygen-rich conditions compared to oxygen-limited environments^17, 47^. In summary, extracellular matrix production in these engineered *E. coli* strains was governed by an interplay between genetics and environmental conditions. While EcN and the curli-producing mutant exhibited measurable ECM production on solid substrates, liquid submersion suppressed both curli and cellulose production to baseline levels regardless of strain genotype.

### Incubation conditions and ECM composition affect bioprint mechanical properties

After identifying variations in ECM production based on bacterial strain and incubation conditions in 3D bioprints, testing was performed to identify any impacts on the mechanical stability of the bioprints. Tensile testing was performed on the incubated, 3D-printed dogbone structures to measure their Young’s modulus (elastic modulus) and ultimate tensile strength (UTS) (Figure 9, Supplementary Figure 5). For all bioprinted structures incubated in liquid LB, no elastic modulus or strength data could be recorded regardless of the bacterial strain (Figure 9B). The samples were too fragile to survive mechanical testing, likely due to substantial water absorption that weakened the material to the extent of structural collapse^48^. This water absorption can be noted by the swelling observed for bioprints incubated in liquid LB relative to bioprints incubated on LB agar (Supplementary Figure 6).

**Figure 9.**
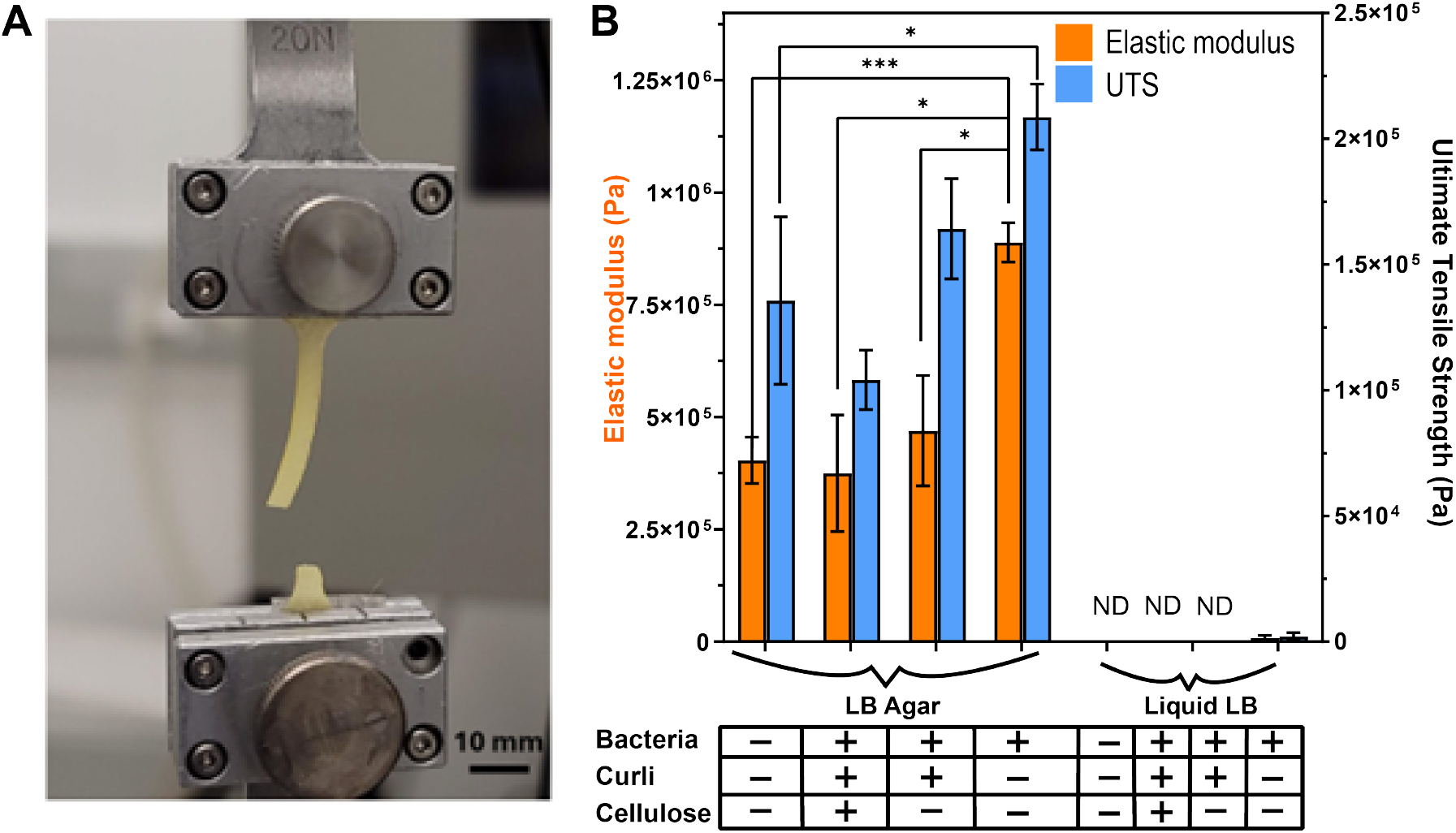
Mechanical integrity of bioprints is governed by extracellular matrix deposition and associated water absorption. (A) Representative photograph of a 3D-printed alginate dog-bone construct mounted in the grips of a universal testing machine after completion of monotonic tensile loading. (B) Elastic modulus (orange bars, left y-axis) and ultimate tensile strength (UTS, blue bars, right y-axis) of alginate-only controls and bioprints embedded with different *E. coli* strains after 7 days incubation on LB-agar or submerged in liquid LB. Error bars were determined using the standard deviation of the samples. Statistical significance was determined using an unpaired Student’s t-test with a Welch’s correction (n=3; * p < 0.05, ** p < 0.01, *** p < 0.001).

In contrast, bioprints that were incubated on the surface of an LB-agar plate, where water uptake was more restricted, showed mechanical characteristics that were quantifiable and varied depending on the strain of *E. coli* that was encapsulated (Figure 9B). Structures with *E. coli ΔcsgA* pGFP-control (curli^−^/cellulose^−^) exhibited a significant increase in both elastic modulus (p = 0.0003 (12.48,3.889)) and ultimate tensile strength (UTS, p = 0.0480 (3.539,2.604)) relative to control alginate prints. This result indicates that bacterial incorporation alone can enhance the physical properties of the alginate hydrogel. However, the ECM producers EcN (curli^+^/cellulose^+^) and *E. coli ΔcsgA* pCsgA-GFP (curli^+^/cellulose^−^), when cultured on agar, consistently exhibited significantly lower elastic modulus and ultimate tensile strength compared to the non-ECM-producing *E. coli ΔcsgA* pGFP (p < 0.05 for all comparisons). These outcomes may be a result of the hygroscopic properties of the ECM components, which have been observed to absorb moisture from their environment to contribute to cellular survival^49^. This excess moisture can be noted by the higher swelling observed for dogbones containing ECM-producing microbes relative to the non-ECM-producing strain (Supplementary Figure 6) and may have functioned as a plasticizer within the hydrogel matrix, diminishing the material’s stiffness and strength^48^.

Among the two ECM producers cultured on agar, bioprints including *E. coli ΔcsgA* pCsgA-GFP (curli^+^/cellulose^−^) exhibited less swelling (Supplementary Figure 6) and significantly higher ultimate tensile strength (p = 0.002) in comparison to those with EcN (curli^+^/cellulose^+^), though the elastic moduli remained comparable between the two ECM-producing strains (p > 0.05) (Figure 9B). These data indicate that the specific composition of the ECM, along with its interaction with water, can have significant impact on mechanical properties. The complex extracellular matrix of EcN, which comprises both curli and a substantial quantity of highly hydrophilic cellulose when cultivated on agar, may have resulted in increased water retention that disrupted the alginate cross-linking network, leading to a more pronounced reduction in tensile strength^48^ in comparison to the mostly curli matrix of *E. coli ΔcsgA* pCsgA-GFP^50^.

### *In situ* incubation within support slurry reveals nutrient-dependent trade-offs

To further explore the relation between incubation conditions and material properties, we investigated the effect of incubating the 3D-printed constructs *in situ* within the agarose support slurry, without the extraction step. This approach mimics a scenario in which the printing medium also serves as the long-term culture environment. Bioprints that contained one of three different *E. coli* strains with varying ECM production phenotypes were compared in two incubation conditions: a nutrient-rich slurry containing LB medium (Slurry/LB) and a nutrient-poor control slurry with no additional nutrients (Slurry/H_2_O). Congo Red absorbance values for all strains remained at or below the background levels of the alginate-only control (Supplementary Figure 7A). In the Slurry/LB condition, the EcN strain produced significantly higher amounts of cellulose compared to the alginate-only control (p < 0.001; Supplementary Figure 7B). The curli-producing strain (*E. coli ΔcsgA* pCsgA-GFP) also exhibited cellulose levels that were significantly elevated compared to the alginate control (p < 0.01) although significantly less than the EcN strain (p < 0.001). The non-ECM producing strain showed no significant cellulose accumulation above the alginate baseline (p > 0.05). In contrast, in the Slurry/H_2_O condition, cellulose content for all strains was below the background levels seen for alginate-only prints. While the lack of curli accumulation during slurry incubation parallels the results seen in liquid culture (Figure 8A), the nutrient-dependent bioproduction of cellulose observed in slurry baths indicates key differences between liquid and slurry conditions.

The mechanical testing of these *in situ* incubated constructs revealed that the constructs incubated in nutrient-poor slurry (Slurry/H_2_O) were significantly stiffer and stronger than those incubated in the nutrient-rich LB slurry (p < 0.05 for surviving prints; Supplementary Figure 9). The non-ECM-producing *E. coli* samples in the nutrient-rich slurry were too fragile to be measured, further highlighting the loss of mechanical integrity. These results are consistent with our previous data showing the impact of ECM on the mechanical properties of the bioprints (Figure 9B). In the Slurry/LB environment, all strains showed similar or lower elastic modulus and UTS compared to the alginate-only control (Supplementary Figure 8A). Overall, these results indicate that nutrient availability during slurry incubation drove extensive ECM production at the cost of mechanical integrity. This data can be interpreted to indicate that in the slurry/LB environment, the bacteria thrived and produced a dense, hydrophilic ECM that absorbs higher amounts of water from the surrounding hydrogel, leading to a softer, weaker final construct^48^. Conversely, in the Slurry/H_2_O environment, the bacteria were less metabolically active, producing minimal ECM. The resulting constructs had lower water content (Supplementary Figure 6) and thus exhibited better mechanical performance^48^. These results emphasize a critical design principle for ELMs: maximizing biological activity (e.g., ECM production) can negatively impact bulk mechanical properties of the material.

## Discussion

This study shows how to construct and test a DIY 3D bioprinting platform that can fabricate complex bacterial structures with improved geometric complexity and dimensional accuracy. A cost-effective, open-source 3D bioprinter was created by modifying a commercial FDM printer. The entire cost of the customized system, including the base printer and all custom components, amounted to approximately $450, representing a fraction of the cost associated with commercial alternatives and supports and helping to make bioprinting technology available to more researchers. The use of slurry-based support bath methods with gelatin and agarose microparticles allowed for the creation of structures with aspect ratios up to 8:1 and feature sizes as small as 260 μm. These support slurries allowed for the fabrication of challenging shapes including overhangs, internal channels, and hollow structures. By optimizing printing settings and adding retraction capability to the bio-ink-extruding syringe pump, off-target bacterial placement between printed structures was reduced by nearly two orders of magnitude. The ability to print up to 100 layers while maintaining precise spatial dimensions is an improvement from previous DIY bioprinting methods, allowing for the printing of complex three-dimensional structures.

In contrast to recent approaches utilizing non-reversible granular supports to bioprint ELMs^34, 36^, the agarose and gelatin slurries used in this study are inexpensive, simple to produce, and use readily available polymers, and the gelatin bath is fully thermoreversible. This feature makes it possible to extract the printed construct easily and non-destructively, which is difficult to do with non-meltable supports, and allowed for the evaluation of the mechanical properties of the engineered ELMs under various post-printing conditions. The methods demonstrated here can be used for either extraction (gelatin slurry) or *in situ* incubation (agarose slurry) of bioprints, according to specific application requirements, to regulate and characterize the spatial organization of designed cells.

Long-term viability studies showed that adding supplemental nutrients to slurry support baths can support bacterial survival for 28 days in printed structures. The capacity to construct intricate, high-aspect-ratio bacterial structures while preserving viability over long durations in nutrient-enriched slurry baths can enable the development of functional living materials characterized by advanced architecture and continuous biological activity. Balasubramanian et al.^20^ highlighted the significance of ECM composition in the durability and biological resilience of 3D-printed *E. coli* biofilms, and our findings broaden this to include the bulk mechanical properties of engineered living materials. Mechanical characterization studies showed that incubation conditions, extracellular matrix composition, and the resulting material properties are interconnected in a complex manner. The availability of oxygen is a key factor in the production of cellulose and its mechanical properties, while the hygroscopic nature of ECM components can promote water absorption and correspondingly a decrease in stiffness and ultimate tensile strength. These results can assist researchers to select the most appropriate conditions for culture post printing so that engineered living materials have the desired mechanical properties. Additional investigation into engineered channel networks within bioprints may allow for enhanced nutrient transport and byproduct removal, resulting in more uniform and predictable material properties. Post-processing strategies such as controlled dehydration or secondary cross-linking could potentially also be applied to mitigate swelling-induced softening.

The hardware design and control software for the 3D bioprinter are both open-source, facilitating the research community to make further modifications and improvements that could accelerate progress in 3D bacterial printing and the creation of engineered living materials. The capabilities of this platform can enable the exploration of bacterial community dynamics in specific three-dimensional environments and support the creation of advanced biomaterials applications in biotechnology, environmental remediation, and tissue engineering and fundamental synthetic biology^51^. Future studies could focus on improving the composition of hydrogels and nutrient delivery methods to enhance bacterial viability within printed structures. The difficulties with liquid diffusion found in static printed constructs during incubation may be reduced by the development of dynamic nutrient delivery systems or the addition of controlled-release mechanisms. By utilizing multiple print heads for multi-material printing, it could be possible to construct symbiotic bacterial communities with improved functional capabilities. This study paves the way for the widespread use of accessible bioprinting technologies and moves the field of engineered living materials forward by demonstrating how to build complex bacterial structures, with applications in many scientific and technological areas.

## Methods

### 3D bioprinter construction and setup

A low-cost, open-source 3D bioprinter was constructed by modifying a commercial Creality Ender 3 fused deposition modeling (FDM) printer. The standard FDM print head was substituted with a customized, lightweight syringe pump assembly inspired by the open-source Poseidon syringe pump system^37^. The syringe pump components were fabricated from polyethylene terephthalate glycol (PETG) using FDM printing to ensure chemical resistance and mechanical durability. The pump design was modified to decrease weight by 29% by 3D-printing with 90% infill. Customized mounting brackets were developed for direct attachment to the printer’s X-axis carriage.

The adapted syringe pump was designed to utilize a 10 mL glass Luer-lock syringe. To accommodate the glass barrel and mitigate mechanical compliance, the syringe was secured by custom-designed adjustable screw clamps. The plunger mechanism was redesigned with a captive slot to securely engage the plunger flange. This modification allowed both accurate extrusion and retraction of the bioink. To achieve sterile printing of Biosafety Level 2 (BSL-2) organisms, the printer was housed within a customized polycarbonate container (ClearView Plastics, Sacramento, CA). The enclosure featured an integrated temperature and humidity monitoring system (ClearView Plastics) and was retrofitted with a 25-watt, 254 nm UVC lamp (BAIMNOCM) for surface sterilization.

### G-code generation and slicing parameters

Computer-aided design (CAD) models were created using SolidWorks. G-code instructions for printing were generated using the Slic3r slicing engine integrated within the Repetier-Host software. Various parameters were optimized for bio-ink extrusion. The nozzle diameter was calibrated at 0.254 mm to be consistent with the 26-gauge dispensing needle. The extrusion multiplier was calibrated to 0.004, resulting in a flow rate of approximately 2 mL/hour at a print speed of 23 mm/s.

### Agarose and gelatin slurry preparation

An agarose slurry was prepared to serve as a support medium. First, a 1% (w/v) agarose gel (UltraPure™ Agarose, Invitrogen, Catalog #16500100) was made by dissolving 2 g of agarose in 200 mL of a 10 mM CaCl_2_ solution within a 500 mL heat-resistant jar. The mixture was heated in a microwave for two 1-minute intervals, with swirling in between, to ensure complete dissolution of the agarose. The solution was then cooled and solidified into a gel puck by placing it in a 4°C refrigerator. The edges of the puck were gently separated from the jar walls with a spatula, and an additional 200 mL of 10 mM CaCl_2_ was added. The gel puck was frozen at -20 °C for 20 minutes. Finally, the frozen puck was blended using a hand blender for 120 seconds to create a fine slurry.

The slurry was then subjected to three wash steps. After the initial blending, the slurry was centrifuged at 3000 g for 2 minutes. The supernatant was discarded, and the microparticles were resuspended in 200 mL of 10 mM CaCl_2_. This centrifugation and washing process was repeated two more times for a total of three washes. For immediate use, the final supernatant was discarded, and the compacted slurry was pipetted into the desired printing container. For storage, the supernatant from the final wash was not discarded; the mixture was vortexed and stored at 4°C for up to three days.

A gelatin-based support slurry was prepared following the same procedure as the agarose slurry, with the primary difference being the use of 4.5% (w/v) gelatin instead of 1% agarose.

### Bacterial strains and culture preparation

Three strains of *Escherichia coli* were used in this study: *E. coli* Nissle 1917 (EcN), a probiotic strain that produces both curli and cellulose (curli^+^/cellulose^+^)^45^; *E. coli ΔcsgA* (pCsgA-GFP), an engineered strain that expresses CsgA (curli) upon induction with rhamnose but does not produce cellulose (curli^+^/cellulose^−^)^20^; and *E. coli ΔcsgA* (pGFP-control), an engineered strain that produces neither curli nor cellulose (curli^−^/cellulose^−^)^20^.

For visualization experiments, such as the retraction study, chromogenic *E. coli* strains were used, including Top10 expressing ‘SPiS Pink’ (iGEM part BBa_K1033923)^42^ or ‘meffBlue’ (iGEM part BBa_K1033901)^41^. Unless otherwise specified, bacterial cultures were grown overnight in Luria-Bertani (LB) medium at 37°C with shaking.

### Bio-ink formulation

A 50 mL bacterial culture was cultivated overnight in Luria-Bertani (LB) broth, with the addition of the appropriate antibiotic if necessary to generate the bio-ink. The cells were concentrated by centrifugation at 1500 g for 5 minutes. The resultant cell pellet was resuspended in 10 mL of sterile 4% (w/v) sodium alginate solution in LB and vortexed for homogeneous mixing, resulting in the final bioink. The bio-ink was centrifuged at 1500 g for 2 minutes to eliminate air bubbles formed during mixing. The degassed bio-ink was subsequently placed into a 10 mL glass Luer-lock syringe for the printing process.

### 3D bioprinting and post-processing

Bioink was extruded by a 26-gauge needle into a Petri dish containing support slurry. The alginate in the bio-ink underwent crosslinking upon interaction with calcium ions in the slurry, thereby solidifying the printed structure. Post-printing, gelatin slurries were heated to 40°C for 20 minutes to liquefy the gelatin, allowing the delicate removal of the free-standing bioprint. Structures fabricated in agarose slurries were retained in the slurry for in-situ incubation.

To evaluate print fidelity and dimensional accuracy, cylindrical shapes were generated with a surrounding single-walled ‘skirt’ structure using slic3r slicing software. This skirt was printed with one layer width and with varying total layer counts (1, 10, 20, 30, 40, or 100 layers). The measurements of layer height and width were quantified on these skirt structures.

For aspect ratio characterization, hollow cylindrical structures with a 5-mm outer diameter were printed to varying heights ranging from 5 mm to 40 mm. To compare slurry-based printing with direct printing onto LB agar, identical structures were printed onto LB-agar plates supplemented with 0.1 M CaCl_2_ to induce crosslinking.

For long-term viability experiments, hollow cylinders with 10-mm outer diameter and 5-mm inner diameter were printed to a standard height of 10 mm.

### Colony-forming unit assay

Viable cell counts were determined by performing a 10-fold serial dilution followed by plating. This general procedure was applied to two distinct experiments.

To assess long-term bacterial survival, *E. coli* Top10 cells expressing spisPink were printed into hollow cylindrical structures in agarose slurry. Prints were monitored over 28 days at 37°C under two conditions: one group received nutrient supplementation with LB medium, and a control group was maintained in a PBS-only buffer. Both supplementation solutions were prepared with 0.1 M CaCl_2_ to prevent decrosslinking of the alginate hydrogel. A volume of 3 mL of the respective solution was added onto the surface of the slurry every 3 days. At designated time points, individual cylindrical constructs were removed from the slurry and dissolved in 1.5 mL of 1M sodium citrate.

To quantify off-target bacterial deposition in the retraction study, samples of the support slurry were extracted from the regions between printed structures using a sterile spatula and weighed to a standardized mass of 0.15 g to ensure consistency across all replicates.

For both applications, the resulting cell suspensions were serially diluted in LB broth. Aliquots of 50 µL from each dilution were plated onto LB-agar plates containing the appropriate antibiotic (chloramphenicol, 25 µg/mL). Plates were incubated at 37°C for 24 hours, after which colonies were counted to determine the colony-forming units per milliliter (CFU/mL).

### Sample preparation for ECM and mechanical characterization

For extracellular matrix (ECM) quantification and mechanical characterization, dog-bone structures were 3D printed with a total length of 58 mm, featuring a gauge length of 27 mm, a gauge width of 4.3 mm, and a thickness of 1 mm. 3D-printed dog-bone structures were extracted from the gelatin slurry and incubated for 7 days at 37°C. Samples were divided into two experimental conditions: surface incubation on LB-agar plates and submerged incubation in liquid LB medium. To prevent depolymerization of the alginate matrix, both the solid agar and liquid media were supplemented with 0.1 M CaCl_2_. For the submerged samples, the liquid medium was refreshed once after 3 days.

### Cellulose quantification

Cellulose content was quantified using a modified standard anthrone assay^52^. All chemicals were sourced from Sigma-Aldrich (St. Louis, MO) unless otherwise noted. After the required incubation time for biomaterial growth, the 3D-printed dogbone structures were dissolved in 3 ml of 1M sodium citrate to break down the alginate matrix. A 1 ml aliquot of the resulting sample was treated with 3 ml of an acetic/nitric acid reagent (80% acetic acid, 15% nitric acid, 5% water) and incubated for 1 hour at 100°C in a water bath on a temperature-controlled hotplate to hydrolyze non-cellulosic polysaccharides. After incubation, the samples were centrifuged at 3000 g for 15 minutes, and the supernatant was discarded. The remaining pellet, containing the cellulose, was washed with deionized water and then dissolved in 10 mL of 67% sulfuric acid. The resulting solution was brought to a final volume of 100 mL using deionized water. 1 ml of this diluted cellulose was then reacted with 10 mL of anthrone reagent (0.2% anthrone in concentrated sulfuric acid), prepared fresh daily. The reaction was incubated in a boiling water bath for 16 minutes. After cooling the tubes to room temperature, the absorbance was measured at 620 nm using a spectrophotometer.

A standard curve was developed with known amounts of microcrystalline cellulose. A stock solution of 100 µg/mL was created by dissolving 50 mg of microcrystalline cellulose in 10 mL of 67% sulfuric acid with mild heating, later diluted to a final volume of 500 mL with deionized water. A series of standards was prepared from this stock by diluting certain volumes to a final volume of 5 mL with deionized water. Each 5 mL standard was subjected to cooling in an ice bath prior to the addition of 10 mL of cold anthrone reagent. The tubes were thoroughly mixed and incubated in a boiling water bath for 16 minutes. After letting it cool, the absorbance of the standards was measured at 620 nm relative to a reagent blank containing 5 mL of deionized water and 10 mL of anthrone reagent. The standard curve was then used for determining the cellulose concentration in the dogbone samples. The assay showed a strong linear correlation between absorbance and cellulose concentration (R^2^ > 0.98). A representative standard curve is shown in Supplementary Figure 4.

### Congo Red quantification

Curli amyloid fiber content was determined using a Congo Red binding assay. After incubation, individual 3D-printed dogbone structures were dissolved in 1.5 mL of 1M sodium citrate in a 24-well plate. A 1% (w/v) Congo Red stock solution was added to each well at a ratio of 0.2 µL per mL of citrate solution. The plate was incubated overnight to allow the dye to bind to the amyloid fibers. Following incubation, the supernatant containing unbound dye was carefully removed. The absorbance of the remaining dye bound to the material in each well was then measured directly at 497 nm using a plate reader. This absorbance value corresponds to the relative amount of curli fibers present in the sample.

### Mechanical characterization of bioprints

The mechanical properties of the 3D-printed constructs were analyzed after a 7-day incubation period. Tensile testing was performed on dog-bone shaped samples, according to ASTM D638-05 guidelines, using a Zwick-Roell Z010 universal testing machine equipped with a 20 N load cell (X-Force HP, Zwick-Roell). To minimize slippage, non-woven fabric was applied to the grip sections of the samples. The specimens were subjected to monotonic tensile loading to failure at a constant crosshead displacement speed of 1.1 mm·s^−1^. Specimen width and thickness were measured optically, and the initial gauge length was measured with calipers. Engineering stress was calculated by dividing the recorded force by the initial cross-sectional area, and engineering strain was calculated as the crosshead displacement divided by the initial gauge length. Prior to testing, the thickness and width of the gauge section were measured for each sample condition to calculate the cross-sectional area (Supplementary Figure 6). The stress-strain data (Supplementary Figures 5 and 8) were used to determine the Young’s modulus (elastic modulus) and ultimate tensile strength (UTS). The stress-strain data was retained only until the drop in stress indicating a broken sample. The maximum stress value before the breakage was determined to be the UTS. The slope of the linear elastic region was determined by using a 3 % window to define individual gradients. The maximum gradient from these individual gradients was then identified, to avoid cell loading noise and plateau effects, and presented as the elastic modulus. Three to four independent samples (n=3-4) were tested for each experimental condition.

### Statistical analysis

Quantitative data are presented as the mean ± standard error of the mean (SEM) from at least three independent experiments. Statistical significance between two groups was determined using a two-tailed Student’s t-test. For comparisons of more than two groups, a one-way analysis of variance (ANOVA) was performed. A p-value of less than 0.05 was considered statistically significant.

## Supporting Information

Hardware design specifications and printing instructions, additional figures displaying agarose slurry preparation, print fidelity comparison between printing within slurry and directly onto agar, incubation setup, anthrone assay standard curve, and ECM and mechanical characterization of *in situ* incubated constructs.

## Acknowledgments

The authors wish to thank the Meyer lab members for advice and discussion about the project. Funding to A.S.M. was provided by the U.S. Department of Energy, Office of Science, Basic Energy Sciences via DE-SC0023354, by the Arnold & Mabel Beckman Foundation, by the National Science Foundation via ITE-2137561 and ITE-2230641, and by the University of Rochester. H.C. was supported by the U.S. Department of Energy, Office of Science, Basic Energy Sciences, under Award # DE-SC0025354.

