## Supplemental Figures for "3D bioprinting of engineered living materials in support slurries for complex free-standing structures"

**Open-Source Hardware and Bioprinting Files**

To facilitate reproducibility and the adoption of this low-cost bioprinting platform, all relevant hardware designs, 3D models, and printing instructions have been deposited in a public GitHub repository available at: https://github.com/r1256159/Slurry-Bioprinting-Hardware/tree/main.

Syringe Pump Hardware and Assembly

The bioprinting extruder is a modified version of the open-source Poseidon syringe pump system. We have provided the specific stereolithography (STL) files required to adapt the pump for a Creality Ender 3 printer and a 10 mL glass syringe. These files include:

pump_body_main.stl: The main structural body of the extruder, modified to reduce weight and accommodate the geometry of the glass syringe barrel.

plunger_carriage_retraction.stl: The customized moving carriage featuring a captive slot design. This component secures the syringe plunger flange, enabling the bi-directional linear motion required for both extrusion and precise retraction of the bio-ink.

For general assembly instructions, wiring diagrams, and bill of materials, researchers are referred to the original Poseidon documentation at https://pachterlab.github.io/poseidon/hardware. Users should substitute the standard Poseidon printed parts with the custom STL files listed above.

Bioprinting files:

Experimental geometries and G-code digital files for the specific structures used to characterize the bioprinter are also available in the repository. These files include both the source 3D models (STL) and the machine code (G-code) used for the prints presented in the figures:

lattice_structure_model.stl and the corresponding lattice_structure_print.gcode: The lattice structure used to demonstrate complex, free-standing architecture (Figure 3, Figure 4).

cylinder_5mm_model.stl: The single-walled cylinder model used for layer resolution analysis and aspect ratio quantification (Figure 6).

cylinder_10mm_model.stl and cylinder_10mm_print.gcode: The cylinder model used for retraction optimization (Figure 5) and long-term viability studies (Figure 7).

cylinder_aspect_ratio_print.gcode: The G-code file used to generate the variable-height cylinders for aspect ratio testing (Figure 6).

tensile_dogbone_model.stl and tensile_dogbone_print.gcode: The dogbone shapes used to fabricate constructs for mechanical characterization (Figure 9).

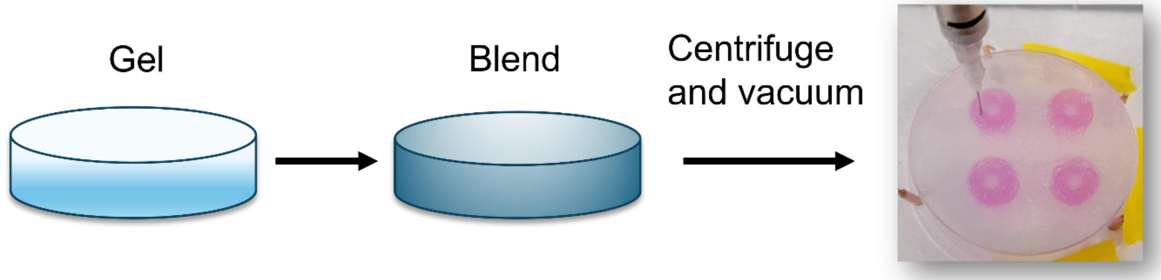

Supplementary Figure 1. Preparation of agarose support slurry. (A) 1% (w/v) agarose is dissolved in a $CaCl_2$ solution and solidified into a puck-shaped gel. (B) The agarose puck is frozen and blended to create a fine microparticle slurry. (C) The slurry is used as a support bath for 3D bioprinting, enabling the fabrication of complex structures.

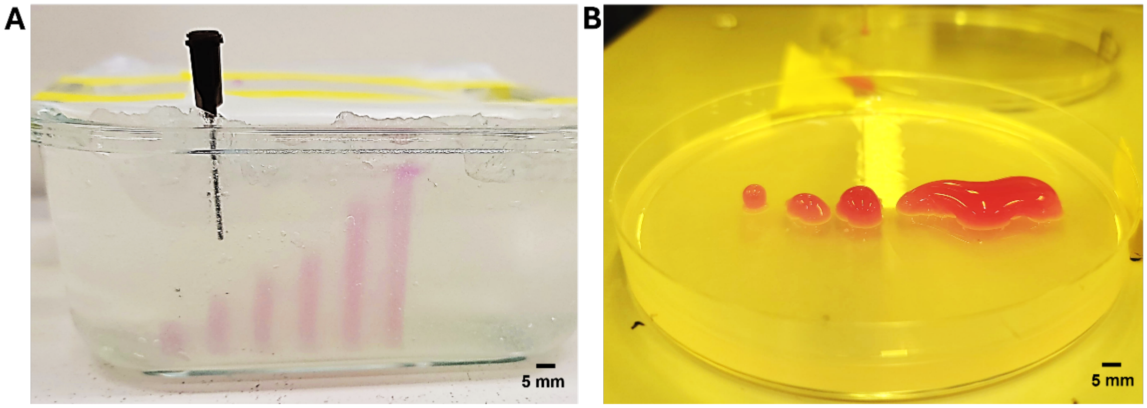

Supplementary Figure 2. Comparison of printing in slurry versus direct printing onto agar. (A) High-aspect-ratio structures printed with *E. coli* expressing the 'SPiS Pink' chromoprotein maintain their vertical form when printed into an agarose support slurry. (B) Identical prints printed directly onto the surface of LB-agar containing $CaCl_2$ collapse under their own weight.

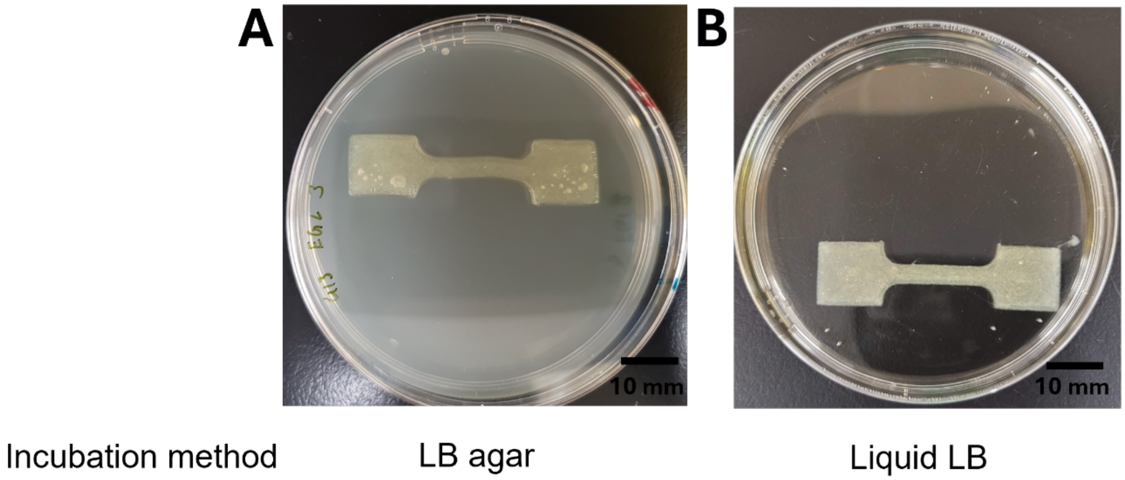

Supplementary Figure 3. Incubation of 3D-printed dog-bone constructs. Representative images of alginate dog-bone structures containing *E. coli*. Constructs were printed within a gelatin support slurry, extracted, and incubated for 7 days either (A) on the surface of an LB-agar plate or (B) fully submerged within liquid LB medium prior to ECM quantification or tensile testing. The images were taken shortly after the transfer from slurry to incubation substrate.

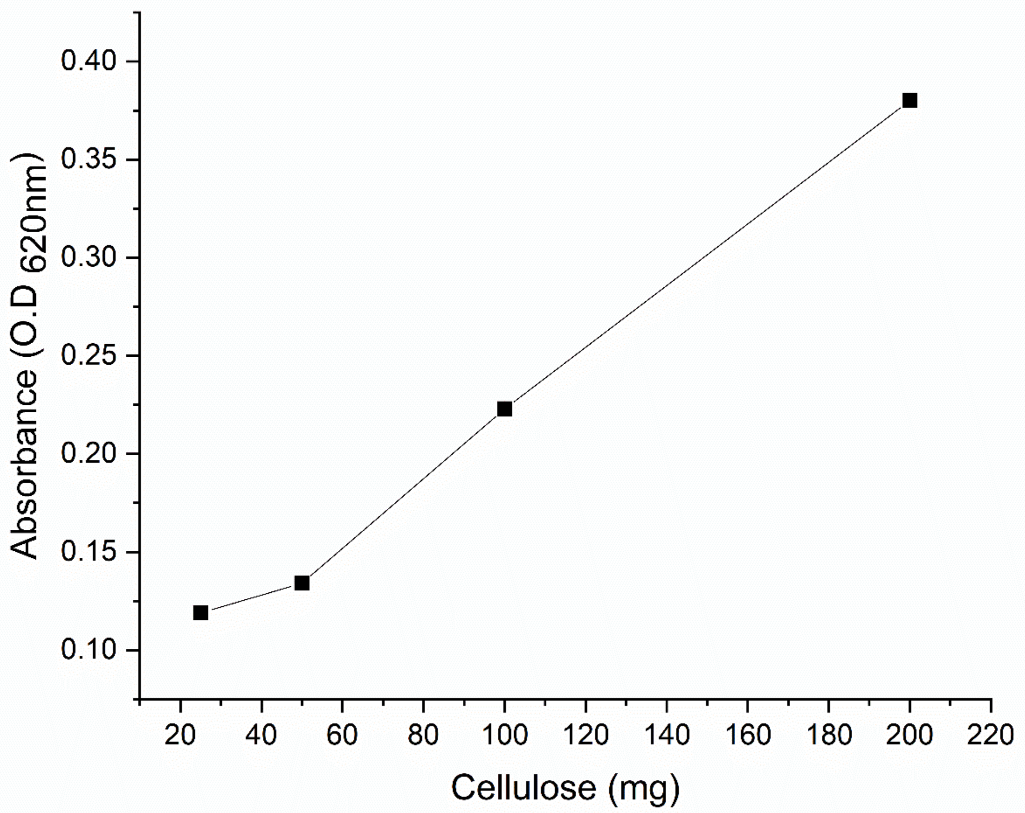

Supplementary Figure 4. Representative standard curve for the anthrone-based cellulose quantification assay. The assay demonstrated a linear relationship ($R^2 = 0.976$) between known concentrations of microcrystalline cellulose and absorbance at 620 nm.

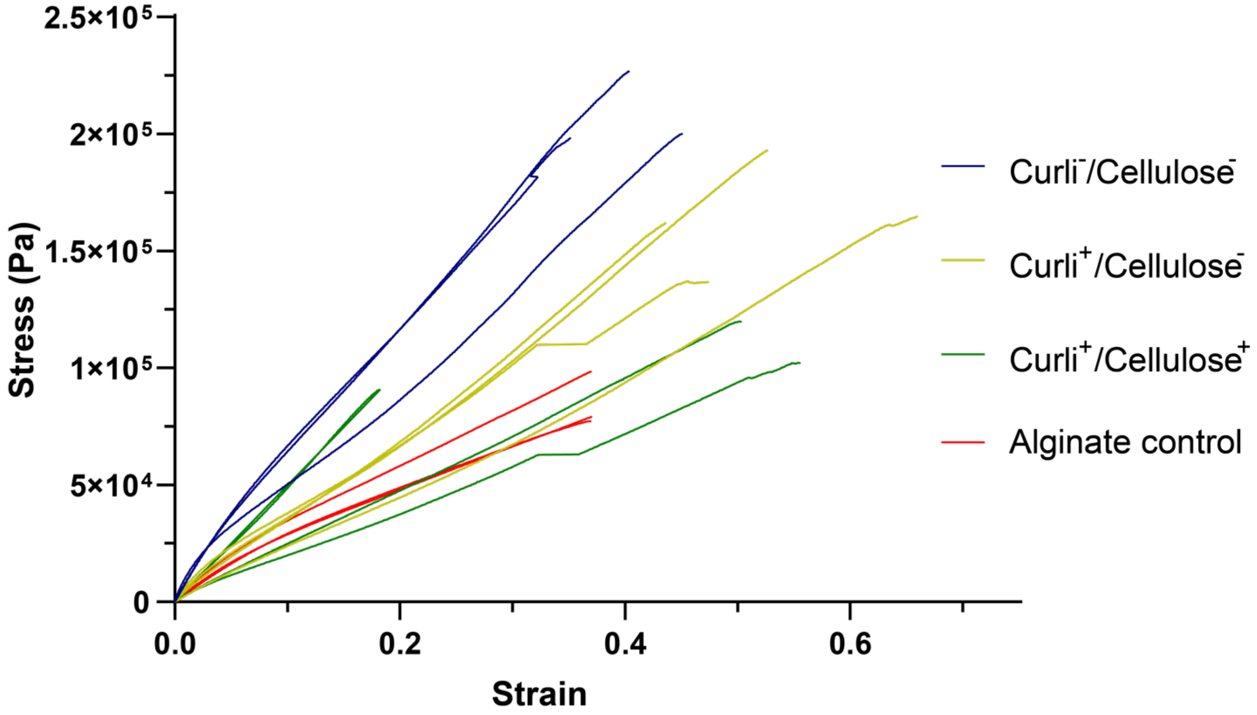

Supplementary Figure 5: Stress-strain behavior of 3D-printed dogbone structures incubated on LB agar. Curves represent n=3-4 independent samples per condition.

| | Incubation Condition | Mean Thickness (mm) | Mean Width (mm) | Mean CSA (mm²) |
|---|---|---|---|---|
| Alginate Control | LB Agar | 0.75 ± 0.11 | 3.13 ± 0.15 | 2.34 ± 0.38 |
| | Slurry/LB | 1.52 ± 0.08 | 4.23 ± 0.06 | 6.58 ± 0.34 |
| | Slurry/$H_2O$ | 2.09 ± 0.19 | 4.24 ± 0.14 | 8.86 ± 1.11 |
| Curli + / Cellulose + | LB Agar | 0.5 | 3.3 | 1.65 |
| | Slurry/LB | 1.50 | 4.00 | 6.00 |
| | Slurry/$H_2O$ | 1.50 | 4.00 | 6.00 |
| Curli + / Cellulose − | LB Agar | 0.5 | 2.75 | 1.375 |
| | Slurry/LB | 1.68 | 4.30 | 7.22 |
| | Slurry/$H_2O$ | 1.50 | 4.20 | 6.30 |
| Curli − / Cellulose − | LB Agar | 0.5 | 2.56 | 1.23 |
| | Liquid LB | 1.6 | 4.19 | 6.7 |
| | Slurry/$H_2O$ | 2 | 4.00 | 8.00 |

Supplementary Figure 6: Dimensions of 3D-printed dogbone constructs after incubation and prior to tensile testing. Dimensions were recorded immediately prior to tensile loading, including thickness, width, and cross-sectional area (CSA). To minimize sample handling and prevent damage that could influence mechanical performance, a single representative construct was measured for each sample containing bacteria. Samples containing curli$^+$/cellulose$^+$ or curli$^+$/cellulose$^-$ bacteria and incubated in liquid LB, and samples containing curli-/cellulose- bacteria and incubated in slurry/LB were too fragile to be measured.

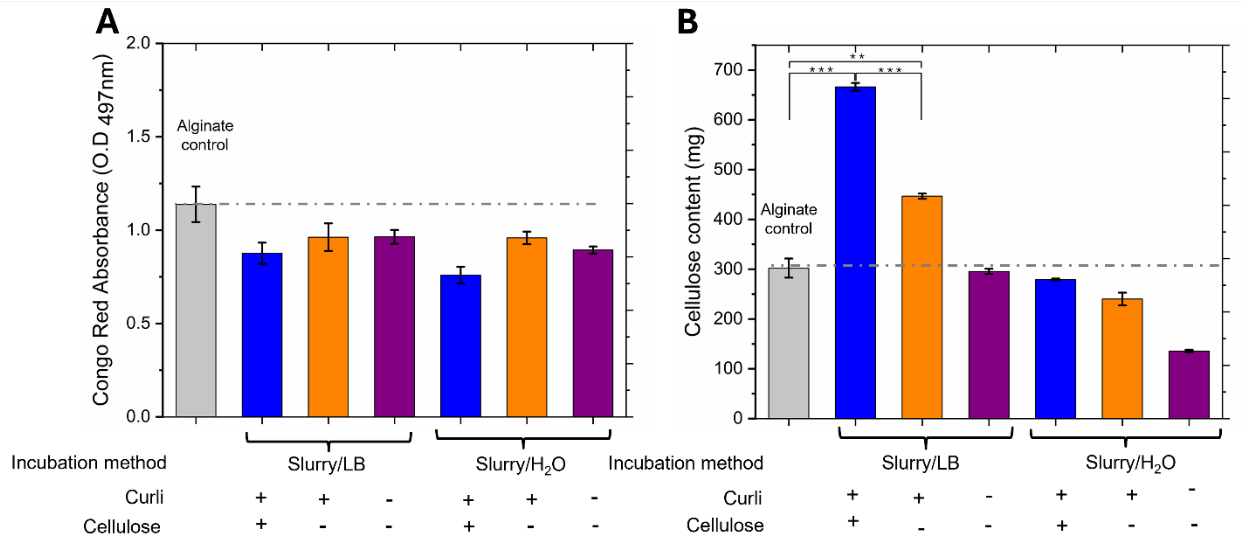

Supplementary Figure 7. ECM production during *in situ* incubation in slurry is nutrient-dependent. (A) Congo Red absorbance and (B) cellulose content for dogbone bioprints containing *E. coli* strains and incubated within either nutrient-rich (Slurry/LB) or nutrient-poor (Slurry/$H_2O$) agarose slurries. Values represent mean ± SEM (n=3). Statistical significance was determined using an independent Student's t-test. No significant differences were observed for Congo Red absorbance values in (A). In (B), asterisks indicate significant differences compared to the alginate control or between indicated groups (** $p < 0.01$, *** $p < 0.001$).

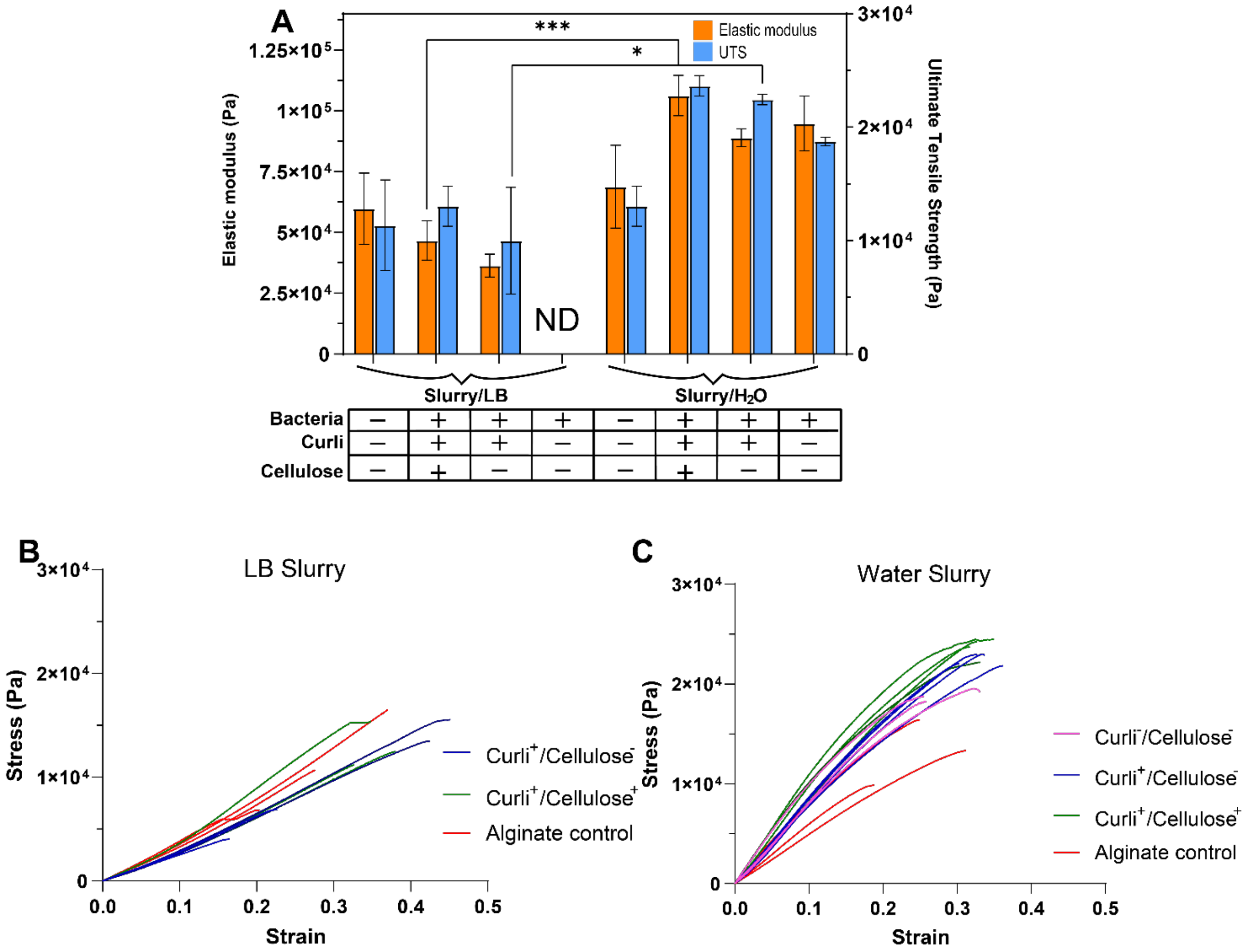

Supplementary Figure 8. Mechanical properties of dogbone constructs incubated *in situ*. (A) Young's modulus (elastic modulus) and ultimate tensile strength (UTS) of constructs incubated within nutrient-rich (Slurry/LB) or nutrient-poor (Slurry/$H_2O$) agarose slurries. Statistical significance was determined using an independent Student's t-test comparing the nutrient-poor (Slurry/$H_2O$) vs. nutrient-rich (Slurry/LB) condition for each strain (* $p < 0.05$, ** $p < 0.01$, ***$p < 0.001$). (B, C) The stress vs. strain curves for samples incubated within (B) the LB-containing slurry and (C) the water-containing slurry.
